# Causal regulators of human T-cell and macrophage activation from perturbation screens, and a replicated coupling of T-cell exhaustion markers with macrophage M1 activation in patients under checkpoint blockade

**DOI:** 10.64898/2026.11.18.767605

**Authors:** Francesca Cupaioli, Ivan Merelli

## Abstract

Engineering anti-tumour immunity requires knowing which genes control T-cell and macrophage programs, and how the two compartments relate to each other in patients. We built ImmunoTwin, an analysis framework that addresses both questions with public data and with every test, baseline and criterion fixed before the result was computed. From a genome-scale CRISPR-interference Perturb-seq in primary human CD4+ T cells (22 million cells, four donors) we derive a map of regulators of the cytotoxic effector program that is calibrated, reverses direction under CRISPR activation (ρ = −0.53), and tracks interferon-γ production among the hits of independent sorting-based screens (ρ = 0.69). From a Perturb-seq in primary human macrophages we recover the TLR4–NF-κB pathway as regulator of the lipopolysaccharide response (pathway-level P = 4 × 10^−4^) and find that transcriptional effects track TNF protein measured in independent donors (ρ = 0.58). In both cell types, however, a generic measure of perturbation strength recovers the hits of functional screens better than the program-specific effect (area under the ROC curve 0.96 versus 0.73 in T cells; 0.87 versus 0.72 in macrophages). In 90 patients from seven cohorts with tumour sampled before and during checkpoint blockade, T-cell and macrophage programs shifted together, most strongly T-cell exhaustion markers and the macrophage M1 program (r = 0.43). Fixed as a hypothesis and tested once in 86 further patients from six cohorts, this association replicated (r = 0.51, permutation P = 10^−4^) and was specific against expression-matched random gene sets (P = 0.004). The baseline state of one compartment did not improve the forecast of the other, so the criterion we had set for calling the framework a patient-level digital twin was not met. ImmunoTwin provides two validated perturbation maps, a replicated cross-compartment association in patients, and a caution on how such maps are benchmarked.

**Author summary:** Immunotherapies work by changing the behaviour of immune cells inside tumours, chiefly T cells, which kill cancer cells, and macrophages, which can either help or hinder them. To design better therapies we need to know which genes switch these behaviours on and off, and whether the two cell types act in concert in patients. We reanalysed large public experiments in which thousands of genes were turned down one at a time in human T cells and macrophages, and built maps of the genes that control the activation of each cell type. The maps agree with independent experiments, but we also found a pitfall: a gene can look important for a specific function because disturbing it upsets the cell broadly. We then examined tumour samples taken from patients before and during immunotherapy. Changes in T cells and in macrophages went hand in hand, and one specific link, between markers of T-cell exhaustion and inflammatory macrophage activity, held up in a second, independent group of patients. However, knowing the state of one cell type before treatment did not help predict the other afterwards. We decided how each analysis would be judged before running it, and we report failed tests alongside successful ones.

## Introduction

Cell therapies and checkpoint antibodies act on a tumour microenvironment in which T-cell function and macrophage state are both decisive [1,2]. Rational design of interventions on either compartment needs two kinds of knowledge: which genes causally control the relevant programs in human cells, and how the two compartments relate to each other in patients under therapy.

Until recently the first question could be approached in primary human immune cells only through co-expression, which does not separate cause from correlation. Pooled CRISPR screens with single-cell read-out (Perturb-seq) [3,4] have changed this. A genome-scale CRISPR-interference Perturb-seq is now available for primary human CD4+ T cells in three stimulation states [5], genome-wide CRISPRi and CRISPRa screens with cytokine read-outs exist for the same cell type [6], and a Perturb-seq with matched FACS screens has been reported for primary human monocyte-derived macrophages [7]. These resources make it possible to ask, with measured effects of real perturbations, which genes move a functional program and whether a transcriptional effect predicts a functional one.

Benchmarks of perturbation-effect prediction have shown that simple baselines are difficult to beat and that an apparently good result can reflect a generic property of the data rather than the biology of interest [8]. An earlier version of this work did not escape the problem: its perturbation models were trained on co-expression-derived pseudo-perturbations, did not exceed co-expression or ridge baselines, and several of its claims weakened or disappeared when controls were added after the fact. We therefore rebuilt the analysis on real perturbation data and adopted one rule throughout: data sources, gene sets, scoring, replication units, baselines, specificity controls and success criteria were written down and time-stamped before the corresponding effects were computed, and departures from that plan are listed (Methods; S1 Text). The framework and its code are called ImmunoTwin, after its original aim — a digital twin of the T-cell– macrophage axis that would forecast an individual patient’s immune state under therapy. Whether that name is deserved was itself made the subject of a pre-specified test.

We report four results. A regulator map of the T-cell effector program is calibrated and consistent in direction with independent gain-of-function data and with functional screens. Recovery of functional screen hits is dominated by generic perturbation strength, to which program-level effects add a small amount of information. A regulator map of the macrophage LPS response recovers the known pathway and tracks TNF production in independent donors. In patients, T-cell and macrophage programs change together under checkpoint blockade; one component of this coupling, between T-cell exhaustion markers and the macrophage M1 program, replicates in independent cohorts, but the baseline state of one compartment does not forecast the other, as a digital twin would require.

## Results

### Pre-specified questions and criteria

Table 1 lists every pre-specified criterion with its outcome (exact values in S1 Table). Briefly, the T-cell arm asked whether real perturbations identify regulators of the effector (GZMB, PRF1, IFNG, GNLY, NKG7, GZMA, FASLG) and exhaustion-marker programs reproducibly across donors (Q1), and whether the effect of a knockdown on the effector program recovers the regulators of interferon-γ and interleukin-2 production found by independent FACS screens, above pre-specified baselines (Q2, primary). The macrophage arm asked the same of the M1 program against a TNF screen, and whether the canonical TLR4–NF-κB pathway is recovered. The patient-level arm asked whether changes in the T-cell effector program and in the macrophage M1 program are coupled within patients under checkpoint blockade (D1), and whether adding the baseline state of the other compartment improves the forecast of a patient’s on-treatment state (D2). The words “digital twin” were to appear in the title only if D2 was met.

**Table 1.** Pre-specified criteria and outcomes.

| Question | Criterion (fixed before the analysis) | Observed | Outcome |
|---|---|---|---|
| Q1. T-cell regulator map | ≥ 20 genes at q < 0.05 on the effector program in ≥ 1 condition | 33 (rest), 4 (8 h), 6 (48 h) | Met |
| Q1. Reproducibility | Leave-one-donor-out effect correlation > 0.5 | 0.15 (all targets); 0.27 (top 5%) | Not met |
| Q1. Calibration | Control pseudo-targets called | 0 of 442 in each condition | Passes |
| Q2. Functional validation (primary; IFN-γ, 8 h) | AUROC ≥ 0.65, CI excluding 0.5 | 0.73 (0.58−0.86); 27 hits, 6,215 genes | Met |
| Q2. Baselines of the original plan | Above target expression, essentiality proxy, random | 0.68, 0.55, 0.52; CI of difference −0.11 to 0.19 | Not met |
| Q2. Strength baseline (added in Addendum A, before any effect) | Above the number of DE genes after knockdown | 0.96; CI of difference −0.37 to −0.10 | Not met; predictor below baseline |
| C1. Beyond strength | Likelihood-ratio P < 0.05 given strength, expression, essentiality | OR 1.45, P = 0.0011 | Met |
| C2. Matched comparison | Hits lower than strength-matched non-hits, P < 0.05 | 70%, P = 0.0065 | Met |
| E2. Macrophage functional validation (TNF screen; M1 without TNF) | AUROC ≥ 0.65, CI excluding 0.5; reported against baselines | 0.72 (0.60−0.84); strength 0.87 | AUROC met; below strength baseline |
| E4. TLR4−NF-κB pathway | Mean effect below random sets, P < 0.05 | −0.28; P = 4 × 10<sup>−4</sup> | Met |
| D1. Coupled changes in patients (effector−M1) | r > 0, permutation P < 0.05 and specificity P < 0.05 | r = 0.23, P = 0.034, specificity P = 0.086; n = 90 | Not met |
| D2. Patient-level forecast | Twin error lower than own-compartment, Holm P < 0.05, ≥ 1 of 2 targets | M1 +0.11 (worse, P = 0.014); effector +0.04 (P = 0.52) | Not met |
| D5. Title | “Digital twin” in the title only if D2 met | — | Not in the title |
| F1. Replication of exhaustion−M1 | r > 0, permutation P < 0.05, ≥ 25 patients not used above; program-specific if specificity P < 0.05 | r = 0.51, P = 10<sup>−4</sup>, specificity P = 0.004; n = 86, six cohorts | Met; program-specific |

### A calibrated, directionally consistent map of effector-program regulators in primary human T cells

The CD4+ T-cell Perturb-seq comprises 21,996,265 cells from four donors with CRISPRi knockdown of 12,731 genes, profiled at rest and 8 h and 48 h after stimulation [5]. For each target and donor we computed the change in the effector program relative to non-targeting controls from pseudobulk profiles, and tested the four donor-level effects with a variance-moderated *t* statistic [9]. Targets without a significant on-target knockdown were excluded, as were the program genes themselves; at rest this left 6,187 of 9,849 targets measured in all four donors.

At rest, 33 knockdowns changed the effector program at a false-discovery rate of 5% (32 decreases, one increase; 28 also specific against expression-matched random gene sets; Fig. 1a; S3 Table). They include the lineage transcription factor *TBX21* (Δ = −1.33, q = 0.038), *STAT5B, BHLHE40* and the T-cell-receptor adaptor *LCP2*. Fewer were found after stimulation (four at 8 h — *LCP2, CD3G, STAT3, SETD1B* — and six at 48 h). The exhaustion-marker program had 22 regulators at 8 h, 21 of them decreasing it and several belonging to proximal T-cell-receptor signalling (*CD3E, CD3G, LAT, LCP2, PLCG1*), consistent with this marker set reporting activation rather than exhaustion in acutely stimulated cells.

**Figure 1.**
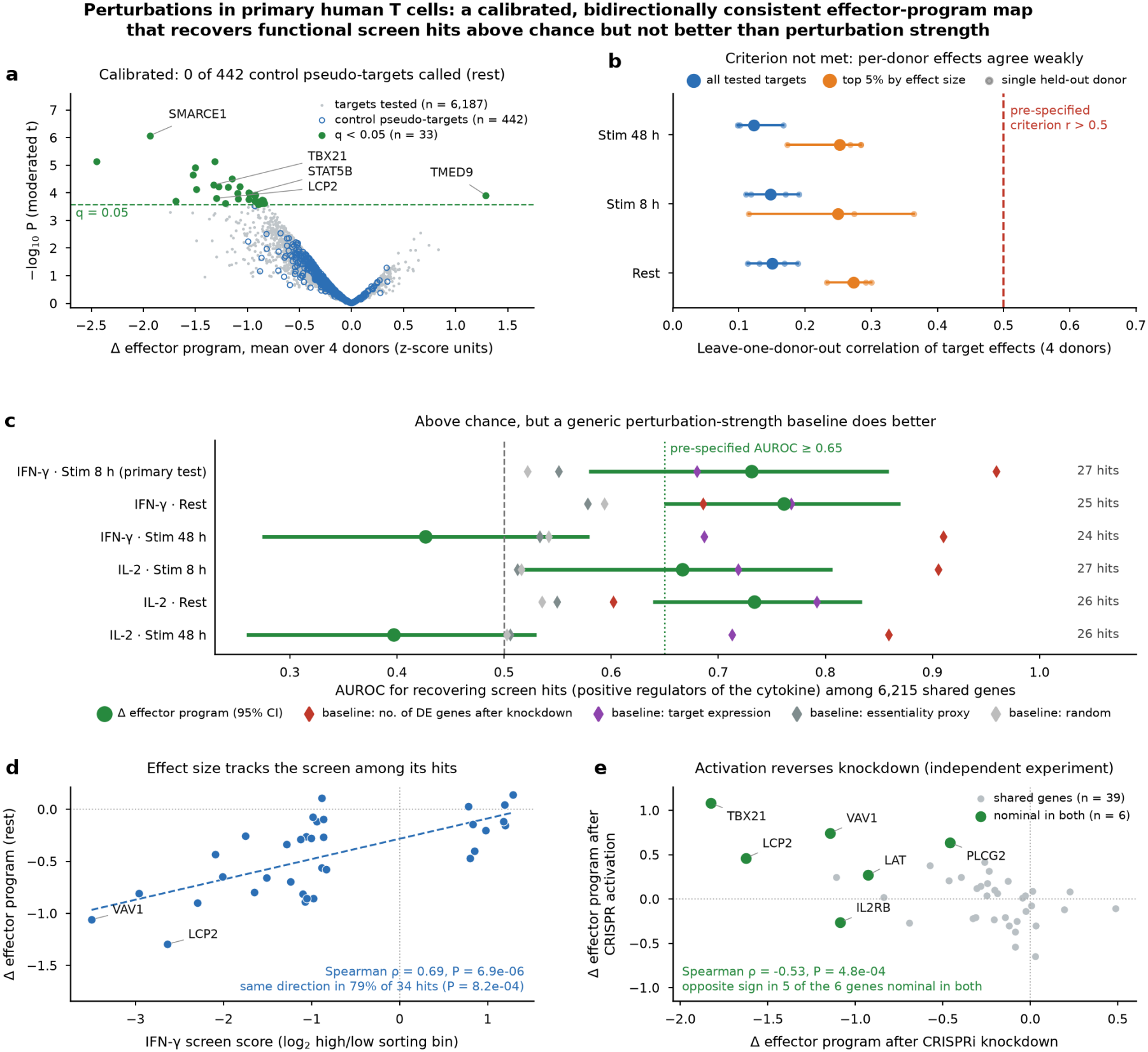
Regulators of the effector program in primary human CD4+ T cells from genome-scale CRISPRi Perturb-seq. (a) Effect of each knockdown on the effector program at rest (mean over four donors) against the moderated-*t* P value; green, q < 0.05; open blue, control pseudo-targets. (b) Leave-one-donor-out correlation of effects, all targets and the 5% largest, against the pre-specified criterion. (c) Recovery of positive regulators of interferon-γ and interleukin-2 from independent FACS screens by the effector-program effect (green, bootstrap 95% CI) and by baselines. (d) Effector-program effect against screen score among screen hits. (e) Effect of CRISPR activation against CRISPRi knockdown of the same genes.

The map is calibrated: pseudo-targets built by pairing non-targeting guides at random and passed through the same pipeline gave no call in any condition (0 of 442 at rest). It is, however, under-powered. With four donors the leave-one-donor-out correlation of effects is 0.15 over all targets and 0.27 over the 5% largest effects, below the pre-specified 0.5 (Fig. 1b), and no target is significant with an unmoderated *t* test. The 33 regulators should be read as a conservative list from a noisy map, not as a complete one.

Two independent data sets support the direction and size of the effects. In a CRISPR-activation Perturb-seq of 73 genes in stimulated T cells [6], over-expression moved the effector program opposite to knockdown (Spearman ρ = −0.53, P = 4.8 × 10^−4^ over 39 shared genes with effective knockdown; opposite sign for five of the six genes nominally significant in both, including *TBX21*, +1.08 on activation and −1.83 on knockdown; Fig. 1e). Among the genes called by the interferon-γ CRISPRi FACS screen [6], the change in the effector program correlated with the screen score (ρ = 0.69, P = 6.9 × 10^−6^, n = 34 at rest, same direction for 79%; ρ = 0.54, P = 5.1 × 10^−4^, n = 37 at 8 h; Fig. 1d).

### Perturbation strength dominates recovery of functional hits; program effects add little

The primary pre-specified test asked whether the effector-program effect of a knockdown ranks the 27 positive regulators of interferon-γ production among 6,215 genes shared with the screen. It does so above chance (AUROC 0.73, 95% CI 0.58–0.86, at 8 h; Fig. 1c), meeting the AUROC criterion, and similarly at rest and for interleukin-2, but not at 48 h. It does not exceed the baselines, so the second half of the criterion is not met. Against the three baselines of the original plan (target expression 0.68, an essentiality proxy 0.55, random 0.52) the difference is not significant (CI of the difference from the best of them −0.11 to 0.19).

The fourth baseline, added to the plan before any effect was computed, changes the reading of this result. The number of differentially expressed genes after a knockdown — a measure of how strongly the perturbation disturbs the cell, with no reference to any program — recovers the screen hits with an AUROC of 0.96, significantly above the program effect (CI of the difference −0.37 to −0.10; Fig. 1c). Screen hits are, to a large extent, strong perturbations (Fig. 2a). A program-specific predictor evaluated without this baseline would have appeared informative for a reason unrelated to the program.

**Figure 2.**
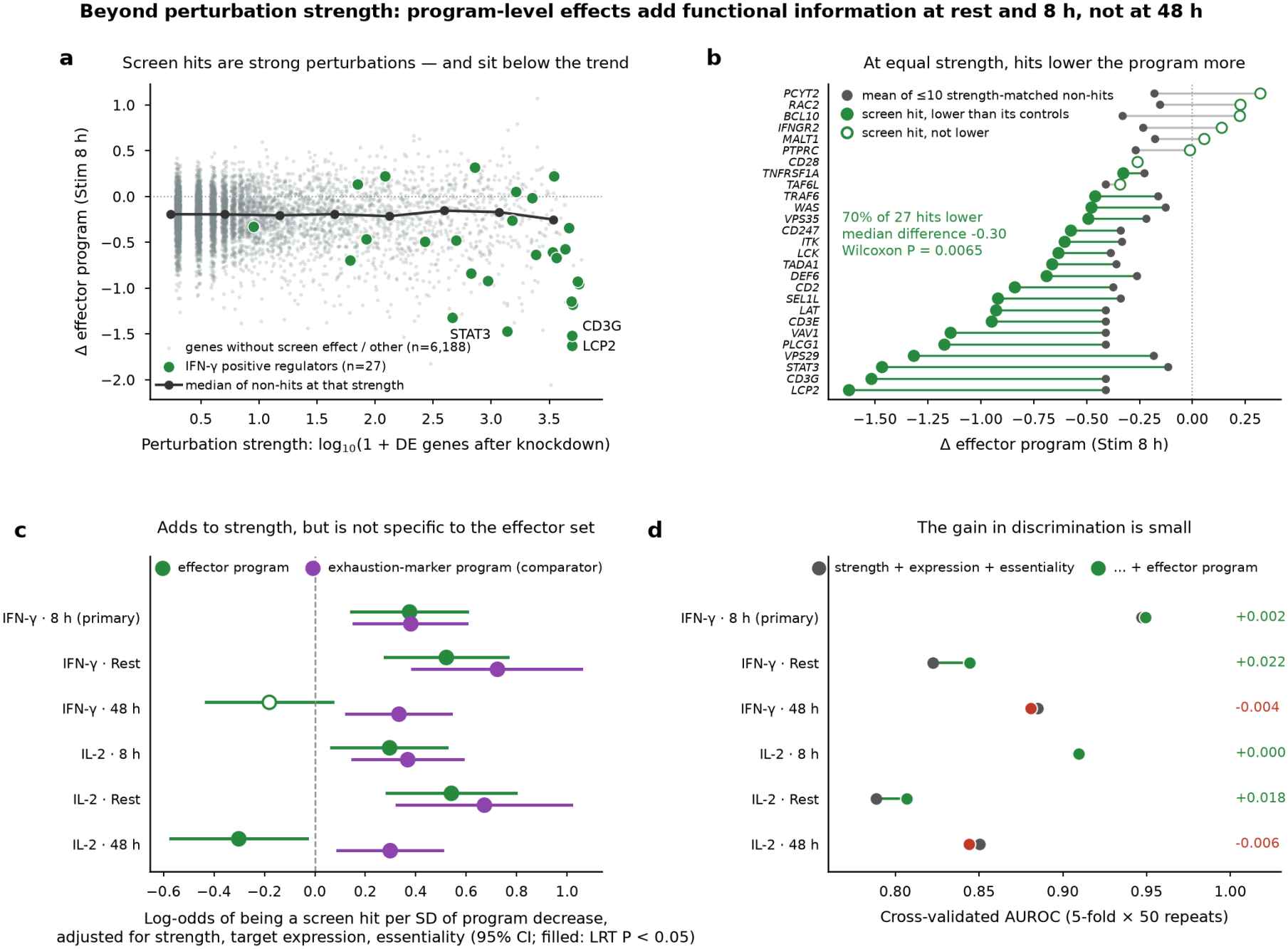
Program-level effects beyond perturbation strength. (a) Effector-program effect against perturbation strength (number of differentially expressed genes) at 8 h; green, positive regulators of inter-feron-γ from the FACS screen; line, median of non-hits. (b) Each screen hit against the mean of up to ten non-hits of matched strength. (c) Coefficient of the program effect in a logistic model of screen-hit status adjusted for strength, target expression and essentiality, for the effector program and for the exhaustion-marker program used as comparator. (d) Cross-validated AUROC without and with the effector effect.

We then asked, with criteria fixed before the analysis, whether the program effect carries information beyond strength. In a logistic model of screen-hit status that includes perturbation strength, target expression and the essentiality proxy, the effector effect is an independent predictor (odds ratio 1.45 per standard deviation, 95% CI of the coefficient 0.14–0.60, likelihood-ratio P = 0.0011), and hits lower the program more than non-hits of matched strength (70% of 27; median difference −0.30; Wilcoxon P = 0.0065; Fig. 2b). Both criteria are met in the primary condition (8 h), and the model gives a similar coefficient at rest (Fig. 2c). Three qualifications apply. The gain in discrimination is negligible (cross-validated AUROC 0.947 to 0.949; Fig. 2d). The effect is not specific to the effector gene set: the exhaustion-marker program gives the same result (odds ratio 1.46, P = 2.9 × 10^−4^; Fig. 2c), the two program effects are correlated across knockdowns (r = 0.41; 0.73 among hits), and neither is clearly informative given the other (post hoc; P = 0.24 and 0.047). And there is no effect at 48 h. What the Perturb-seq adds to perturbation strength is therefore a read-out of lost T-cell activation, not of a particular effector program.

### A regulator map of the macrophage LPS response recovers the TLR4 pathway and tracks TNF production

The macrophage Perturb-seq [7] targets 583 genes with 2,925 guides in primary human monocyte-derived macrophages of one donor, untreated or stimulated with LPS for 5 h. Because all cells come from one donor, guides are the unit of replication and results are descriptive with respect to people. Only 24% of cells carry a guide that accounts for at least 80% of guide reads. A relaxed rule (at least 50%) was adopted after checking, on on-target knockdown alone, that it loses less than 10% of the knockdown (median log_2_ fold change −0.59 against −0.65) while assigning 795,406 instead of 295,770 cells (Fig. 3a); every regulator found under the strict rule keeps its sign under the relaxed one (Fig. 3g).

**Figure 3.**
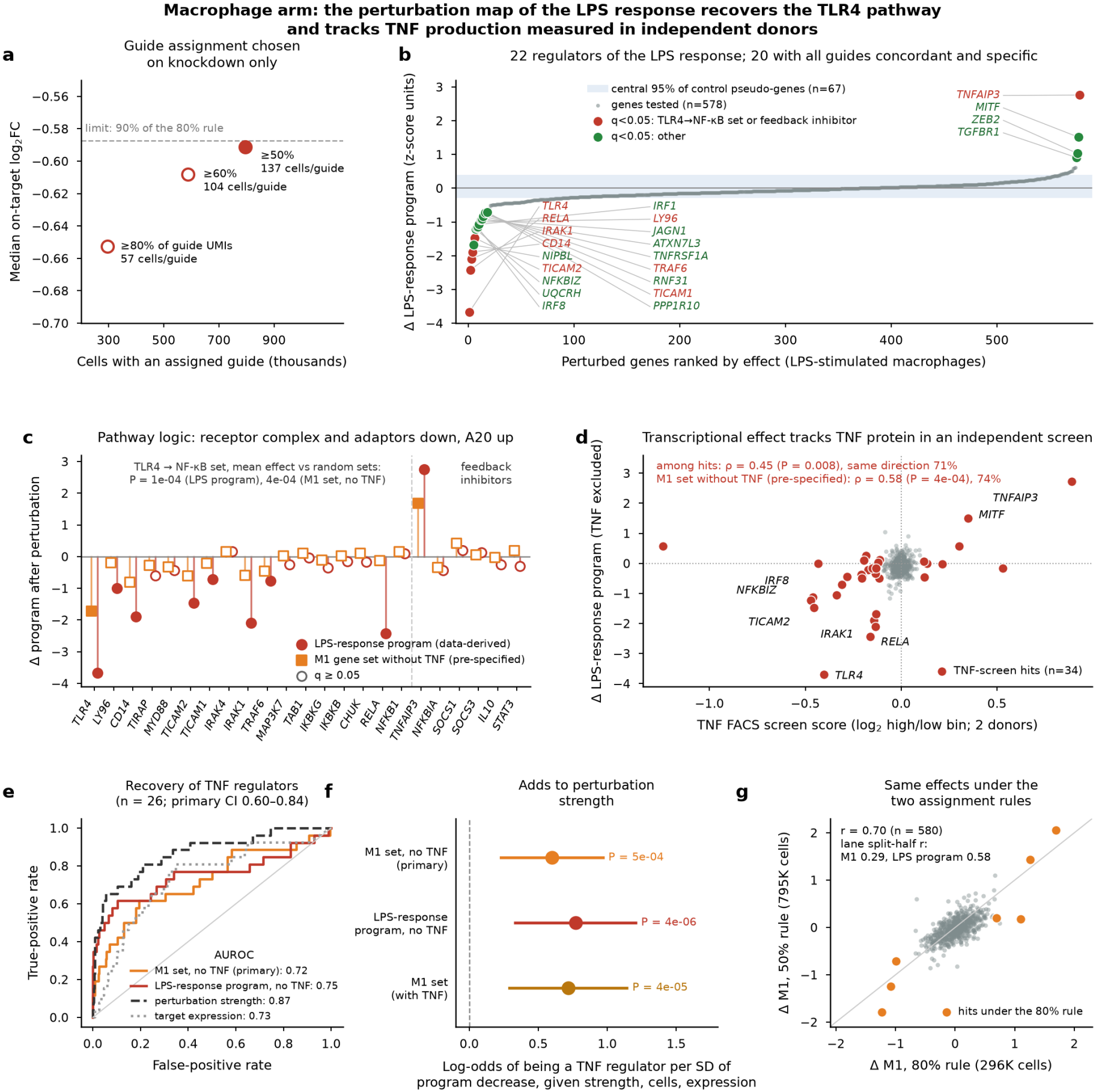
Regulators of the LPS response in primary human macrophages. (a) Cells with an assigned guide against median on-target knockdown for three assignment rules; filled, rule adopted. (b) Perturbed genes ranked by effect on the LPS-response program; band, central 95% of control pseudo-genes. (c) Effects of TLR4–NF-κB pathway members and feedback inhibitors on the LPS-response program and on the pre-specified M1 set without *TNF*. (d) Effect on the LPS-response program (without *TNF*) against the score of an independent TNF FACS screen. (e) Recovery of TNF regulators by program effects and by baselines. (f) Coefficient of the program effect given perturbation strength, cell number and target expression. (g) Effects under the two guide-assignment rules.

The pre-specified M1 gene set turned out to be a poor read-out in these cells: *NOS2, IL12A* and *IL12B* are essentially not expressed, and only *TLR4* and *TNFAIP3* (A20) were called as its regulators under LPS. We therefore also derived, from cells carrying control guides, the 50 genes most induced by LPS and analysed this LPS-response program in the same way; this program was defined after seeing the stimulation check and is labelled post hoc. It has 22 regulators at a false-discovery rate of 5%, 20 of them with all guides concordant and specific against random gene sets (Fig. 3b; S4 Table). They trace the pathway from receptor to transcription factor — *TLR4, LY96, CD14*, the adaptors *TICAM1* and *TICAM2, IRAK1, TRAF6, RNF31, RELA, NFKBIZ, IRF1, IRF8* — with the feedback inhibitor *TNFAIP3* as the strongest gene whose loss increases the response (Fig. 3c). The pathway-level test, fixed in advance on a textbook list of 17 TLR4–NF-κB genes and on the pre-specified M1 set without *TNF*, is met (mean effect −0.28 against −0.02 for the library; P = 4 × 10^−4^ against random sets of the same size; P = 10^−4^ for the LPS-response program). Lane split-half correlation of effects is 0.58 for the LPS-response program and 0.29 for the M1 set.

As functional ground truth we used a FACS screen for TNF protein with the same library in two further donors [7] (34 hits, 26 of them positive regulators; correlation between donors 0.81). The effect of a perturbation on the M1 program computed without *TNF* recovers the positive regulators with an AUROC of 0.72 (95% CI 0.60–0.84; Fig. 3e), meeting the pre-specified criterion, and among screen hits it correlates with the screen score (ρ = 0.58, P = 3.7 × 10^−4^, same direction for 74%; Fig. 3d). The pattern seen in T cells repeats: perturbation strength alone reaches 0.87, the program effect is an independent predictor given strength (likelihood-ratio P = 4.9 × 10^−4^; Fig. 3f) and does not improve discrimination. A second screen, for CD80, had three hits and between-donor correlations of 0.06–0.31, and is not informative.

### In patients, T-cell and macrophage programs shift together under checkpoint blockade, without program specificity or forecasting value

To test the framework at the level of the individual patient we assembled public single-cell cohorts with tumour sampled before and during or after a regimen containing a PD-1, PD-L1 or CTLA-4 antibody, using an inclusion rule applied to deposited metadata before any score was computed. Nine cohorts were prepared [10–18]; seven had at least five patients with at least ten T cells and ten monocytes/macrophages at both time points, giving 90 patients (oesophageal 29, colorectal 18, triple-negative breast 16, head and neck 10 and 6, basal-cell 6, liver 5; Fig. 4a). A breast cohort used in the earlier version of this work [11] is contained in a later release from the same group [10] and was superseded by it.

**Figure 4.**
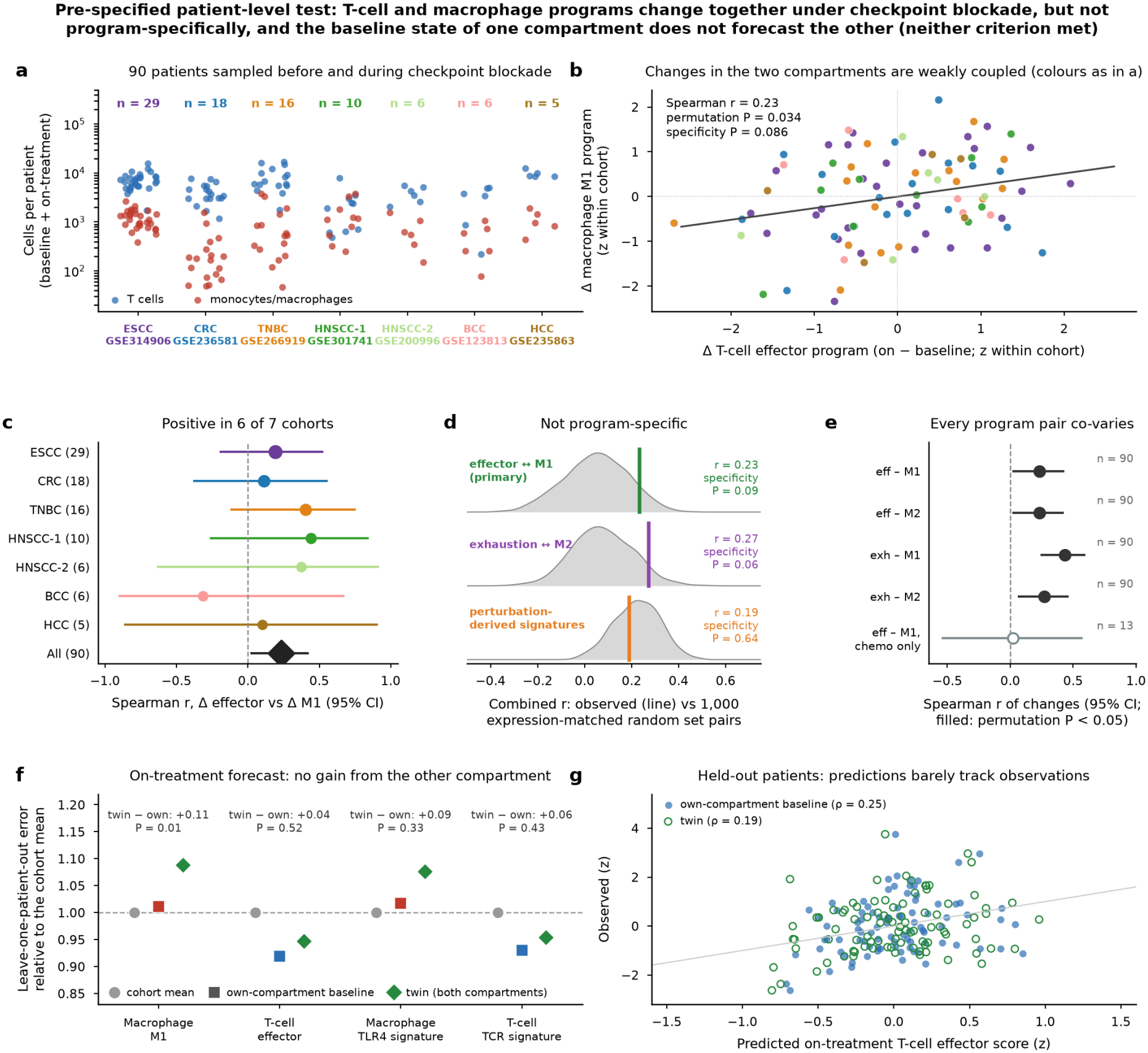
Patient-level test in 90 patients with tumour sampled before and during checkpoint blockade. (a) Cells per patient and compartment in the seven cohorts. (b) Change in the T-cell effector program against change in the macrophage M1 program, standardized within cohort. (c) Correlation per cohort and combined. (d) Observed correlation against expression-matched random gene-set pairs. (e) Correlation of changes for all program pairs and for the chemotherapy-only arm. (f) Leave-one-patient-out forecast error of the on-treatment state relative to the cohort mean, for the own-compartment and twin models. (g) Predicted against observed on-treatment effector score in held-out patients.

#### Coupling of changes (D1)

The change in the T-cell effector program and the change in the macrophage M1 program between baseline and treatment are positively correlated (Spearman r = 0.23 after standardization within cohort; within-cohort permutation P = 0.034; positive in six of seven cohorts; Fig. 4b,c). The pre-specified specificity control is not passed: pairs of expression-matched random gene sets reach r = 0.26 at their 95th percentile (specificity P = 0.086; Fig. 4d), so the criterion is not met. Requiring 30 cells per compartment gives the same picture (r = 0.26, P = 0.035, specificity P = 0.067, n = 68). In 13 patients treated with chemotherapy alone there is no correlation (r = 0.02).

The reason is that the association is not confined to the hypothesised pair (Fig. 4e). Changes in all four pairs of programs are positively correlated, most strongly T-cell exhaustion markers with macrophage M1 (r = 0.43), and signatures derived from the perturbation maps do no better (r = 0.19, specificity P = 0.64). On average both the exhaustion-marker program and the M1 program decrease on treatment (−0.29 and −0.41 log_2_ units; P = 0.014 and 0.002), whereas the effector and M2 programs do not change. In exploratory analyses added after these results, the exhaustion–M1 pair does pass the specificity control, for changes (specificity P = 0.002) and at each time point (baseline r = 0.36, on treatment r = 0.53; specificity P = 0.005 and 0.002). Because this pair was selected after seeing all four, it is a hypothesis; its test on patients not used here is described in the next section.

#### Forecast (D2)

For each patient we predicted the on-treatment score of a program from the baseline score of the same compartment, or from that plus the baseline scores of the other compartment (the twin model), leaving the patient out. Adding the other compartment does not reduce the error: for the M1 program it increases it (difference in squared error +0.11, 95% CI 0.02–0.21, P = 0.014), and for the effector program it makes no difference (+0.04, CI −0.07 to 0.15, P = 0.52); leave-one-cohort-out and the perturbation-derived signatures give the same answer (Fig. 4f,g). Baseline scores themselves predict little: the own-compartment model does not significantly improve on the cohort mean for either program. Baseline programs did not predict clinical response either (79 patients; AUROC 0.32 for T-cell programs and 0.35 with macrophage programs added; descriptive).

Neither criterion being met, the rule fixed in advance applies: the framework is not a patient-level digital twin, and the term does not appear in the title.

### The exhaustion–M1 association replicates in independent patients

Because the exhaustion–M1 pair was selected after seeing the discovery data, we fixed it as a hypothesis, with its test and criterion, before opening any further cohort (Addendum F of the plan): a positive correlation of changes with permutation P < 0.05 in patients not used above, to be called program-specific only if the random-gene-set control is also passed, and a verdict only with at least 25 paired patients. The recorded GEO searches returned 30 further series; seven met the inclusion rule on their deposited metadata [19–22] (three of them, GSE189926, GSE205506 and GSE264586, have no associated publication) and 23 were excluded for reasons listed in the plan. Six cohorts had at least five paired patients — triple-negative breast (31), head and neck (27 and 8), colorectal (8), gastric (7) and renal (5), 86 patients in all — and the seventh, with four, dropped out (S2 Table).

The test was run once. Changes in the T-cell exhaustion-marker program and in the macrophage M1 program were correlated (Spearman r = 0.51, permutation P = 10^−4^; positive in five of six cohorts; Fig. 5a,b), and the correlation exceeded that of expression-matched random gene-set pairs (95th percentile 0.39; specificity P = 0.004; Fig. 5c). Requiring 30 cells per compartment gave the same result (r = 0.55, P = 10^−4^, specificity P = 0.004, n = 82). By the rule fixed in advance, the association is replicated and program-specific. In a post hoc check, the three cohorts in which compartments follow the authors’ own annotation, and therefore do not depend on our cell typing, gave r = 0.50 (P = 10^−4^, n = 66); the three typed here gave r = 0.45 (P = 0.06, n = 20).

**Figure 5.**
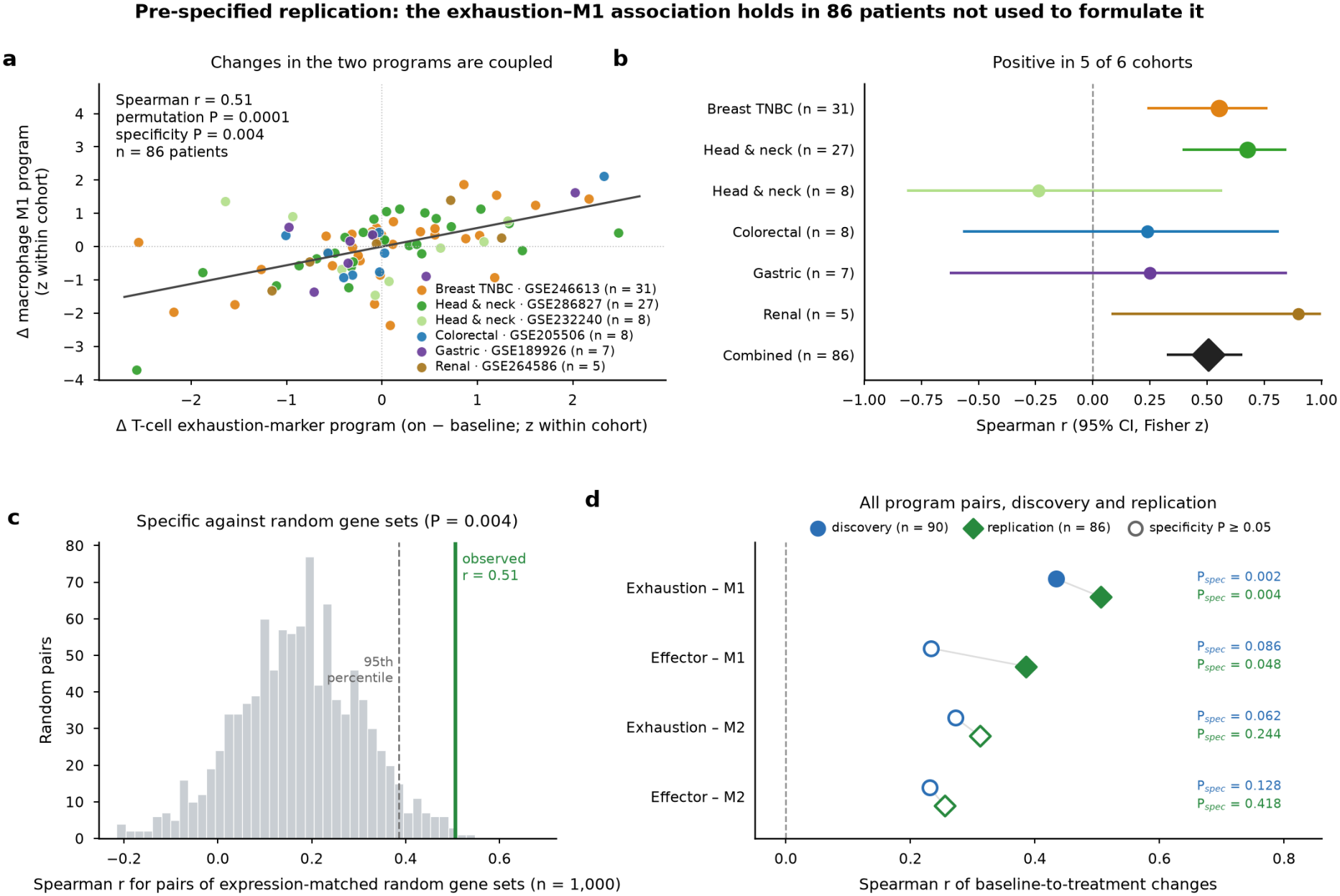
Replication of the exhaustion–M1 association in 86 patients from six cohorts not used in the discovery analysis. (a) Change in the T-cell exhaustion-marker program against change in the macrophage M1 program, standardized within cohort. (b) Correlation per cohort and combined, with 95% confidence intervals. (c) Observed correlation against 1,000 pairs of expression-matched random gene sets. (d) Correlation of changes for the four program pairs in the discovery and replication patients; filled symbols pass the specificity control (P < 0.05). The exhaustion–M1 pair was chosen in the discovery set and tested in the replication set under a pre-specified criterion; the other pairs are secondary.

Secondary analyses, which carried no criterion, complete the picture (Fig. 5d; S5 Table). The effector–M1 pair — the original hypothesis of the patient-level test — was also correlated in the replication patients (r = 0.39, P = 5 × 10^−4^) and passed the specificity control narrowly (P = 0.048); the two pairs involving the M2 program did not pass it (exhaustion–M2 r = 0.31, specificity P = 0.24; effector–M2 r = 0.25, P = 0.42). Random gene-set pairs correlate at 0.17 on average, which measures the general co-variation of the two compartments on which the exhaustion–M1 association stands out. The direction of the average change differed between the two patient sets: in the replication cohorts exhaustion markers increased on treatment (+0.20 log_2_ units, P = 0.005) and the M1 program did not change, whereas both decreased in the discovery cohorts. The association therefore concerns how patients differ from one another in their change, not a common direction of change. Discovery and replication patients were not pooled.

## Discussion

Three conclusions follow, in decreasing order of confidence.

First, public perturbation data now support regulator maps of functional programs in primary human T cells and macrophages that pass controls which co-expression analyses cannot be subjected to: calibration on pseudo-targets, reversal of direction under gain of function, recovery of a textbook pathway, and agreement with protein-level screens in other donors. The maps are small — 33 regulators of the effector program at rest, 22 of the LPS response — because the criteria are conservative and the data are noisier than their size suggests: four donors do not reproduce single-target effects in T cells, and in macrophages a quarter to two thirds of the cells can be assigned to a guide.

Second, the benchmark most often used to validate such maps — recovery of the hits of a functional screen — is dominated in both compartments by how strongly a perturbation disturbs the transcriptome. A count of differentially expressed genes, which knows nothing about cytokines, outperforms the program-specific predictor by a wide margin. Program effects do carry additional information, but in T cells that information is shared between an effector and an exhaustion-marker gene set and is best described as loss of activation. We suggest that perturbation strength be included as a baseline whenever a transcriptional read-out is validated against a functional screen, alongside the linear and co-expression baselines already recommended for expression prediction [8].

Third, in patients the two compartments are coupled, and one component of this coupling is specific and reproducible. Programs of T cells and macrophages move together between baseline and treatment whatever gene sets are used, and on top of this general co-variation the change in T-cell exhaustion markers tracks the change in the macrophage M1 program: the pair emerged from 90 patients, was fixed as a hypothesis, and held in 86 others. The exhaustion-marker set consists of inhibitory receptors induced by T-cell-receptor stimulation and behaves as an activation read-out in the perturbation data; the M1 set includes the interferon-inducible chemokines CXCL9 and CXCL10. The association is thus consistent with the interferon-γ circuit between antigen-experienced T cells and macrophages [23], but the data are correlative and test neither direction of causation. The pair we had originally hypothesised, effector–M1, failed its specificity control in the discovery patients and passed it narrowly, as a secondary analysis, in the replication patients; we do not claim it. And knowing one compartment at baseline does not help to forecast the other on treatment. The framework therefore describes the axis, with one replicated component; it does not predict it.

The failed forecast decides what ImmunoTwin currently is. It was conceived as a digital twin, and we set in advance the test that would justify the name; the test failed. We keep the name for the framework and its repository because the perturbation maps and the patient-level machinery are the components such a twin would need, but we make no claim that it forecasts a patient. Two kinds of data would change this: perturbations in co-culture, where the effect of an edit in one cell type on the other is measured rather than inferred — both Perturb-seq data sets used here are monocultures — and longitudinal cohorts large enough to estimate patient-level effects.

### Limitations

The T-cell data are CD4+ cells stimulated in vitro, whereas tumour exhaustion is mainly a CD8+ phenotype of chronic stimulation; the exhaustion-marker set behaves as an activation read-out in these data. The macrophage Perturb-seq is a single donor and a single stimulus; the LPS-response program was defined after seeing the data. CRISPRi knockdown is partial. The patient cohorts differ in tumour type, regimen, platform and timing of the on-treatment sample; standardization within cohort removes level differences but not heterogeneity of effect, and some regimens include chemotherapy. Program scores are means over small gene sets in pseudobulks of as few as ten cells. Nothing in this work measures tumour killing or tumour control.

## Methods

### Pre-specification

The analysis plan (S1 Text) was written on 13 November 2026 before any effect in the perturbation data was computed, and extended by dated addenda, each stating what had already been seen. It fixes data sets and replication units, the four program gene sets, pseudobulk scoring, tests, multiple-testing correction (Benjamini–Hochberg, q < 0.05), the specificity control against expression-matched random gene sets, baselines and success criteria.

### Gene programs

Effector: GZMB, PRF1, IFNG, GNLY, NKG7, GZMA, FASLG. Exhaustion markers: PDCD1, HAVCR2, LAG3, TOX, ENTPD1, LAYN, TIGIT, CTLA4. M1: TNF, IL1B, IL12A, IL12B, NOS2, CXCL9, CXCL10, CD80. M2: CD163, MRC1, TGFB1, IL10, VEGFA, SPP1, MSR1, MAF. These sets are unchanged from the earlier version of this work.

#### T-cell regulator map

Guide × donor × condition pseudobulk counts of the genome-scale CRISPRi Perturb-seq [5], filtered with the quality flags supplied by its authors, were summed over the guides of each target within donor and condition and converted to log_2_(CPM + 1); non-targeting guides were pooled into one control profile per donor and condition. Within a condition each gene was standardized over all target × donor profiles; the program score is the mean standardized expression of the program genes, and the effect of a target in a donor is its score minus that of the control profile of the same donor. Targets present in all four donors, with a significant on-target knockdown in the table supplied with the data set, and not belonging to the program were tested with a one-sample moderated *t* statistic (empirical-Bayes shrinkage of the variance across targets [9]) and Benjamini–Hochberg correction. Calibration: non-targeting guides were paired at random into pseudo-targets and analysed identically. Specificity: 1,000 random gene sets matched to the program genes by expression bin. Reproducibility: correlation of the effect in one donor with the mean of the other three.

### Functional validation in T cells

Gene-level scores of the genome-wide CRISPRi and CRISPRa FACS screens for interferon-γ and interleukin-2 [6] were computed from guide counts of two donors: per guide and donor, log_2_ of the CPM-normalized ratio (high + 1)/(low + 1) between sorting bins, averaged over donors and then over the guides of a gene; gene-level P values come from an empirical null of 10,000 random guide sets of the same size, with Benjamini–Hochberg correction. Positive regulators are genes whose knockdown lowers the cytokine at q < 0.05. Predictor: the negative effector-program effect (donor mean). Baselines: expression of the target gene in control cells, an essentiality proxy (depletion of the guides in the unsorted sample relative to the plasmid library of the same screen) and a random ranking, listed in the original plan; and the number of differentially expressed genes after knockdown in the statistics supplied with the data set, added in Addendum A before any effect was computed. The Perturb-seq and the screens come from the same laboratory; they are independent experiments with different donors, libraries and read-outs. AUROC confidence intervals and differences use 2,000 bootstrap resamples of genes, with predictor and baselines evaluated on the same resample. The direction check uses the CRISPRa Perturb-seq of ref. 6 scored in the same way.

### Beyond perturbation strength

Logistic regression of screen-hit status on perturbation strength (log_10_ of one plus the number of differentially expressed genes), target expression and the essentiality proxy, with and without the standardized program effect; likelihood-ratio test; 5-fold cross-validation repeated 50 times. Matched comparison: each hit against up to ten non-hits nearest in strength, Wilcoxon signed-rank test.

### Macrophage regulator map

Filtered feature–barcode matrices of the macrophage Perturb-seq (GSE327644 [7]) were read for the four lanes of each condition. A cell is assigned to a guide when it has at least three guide reads of which one guide accounts for a threshold fraction; thresholds of 80%, 60% and 50% were compared on on-target knockdown only, and the most permissive one retaining at least 90% of the knockdown of the 80% rule was adopted. Counts were summed per guide over lanes; guides with at least 20 cells were kept. Program scores and gene-level effects (mean over guides) were computed as above against half of the non-targeting guides, the other half forming pseudo-genes for calibration; P values come from an empirical null of non-targeting guide sets of the same size. The on-target knockdown filter of the plan could not be applied because no efficiency table is supplied and a nuclease knockout does not reliably lower the transcript; this is a deviation. The LPS-response program is the 50 genes with the largest LPS-versus-untreated log_2_ fold change in cells carrying control guides (at least 10 CPM under LPS). TNF and CD80 FACS screens (GSE327124) were scored as for T cells. Pathway test: mean effect of TLR4, LY96, CD14, MYD88, TIRAP, TICAM1, TICAM2, IRAK4, IRAK1, TRAF6, MAP3K7, TAB1, IKBKB, IKBKG, CHUK, RELA and NFKB1 against 10,000 random gene sets of the same size from the library, one-sided.

### Patient cohorts

GEO was searched with recorded queries for single-cell RNA-seq of human tumours sampled before and during or after checkpoint blockade. A series was included if its deposited metadata showed tumour tissue profiled without sorting that removes T or myeloid cells, samples of the same patients before and during or after a regimen containing a PD-1, PD-L1 or CTLA-4 antibody, and processed counts with a patient and time-point label. Included: GSE266919 [10], GSE169246 [11], GSE123813 [12], GSE120575 [13], GSE236581 [14], GSE314906 [15], GSE235863 [16], GSE200996 [17], GSE301741 [18]. Compartments follow the authors’ annotation (T cells; monocytes and macrophages), with a marker-based rule inside the myeloid class where sub-labels are unnamed; where no annotation is deposited, Leiden clusters were typed with immune marker sets. Counts were summed per sample and compartment, pooling libraries or tumour regions of the same patient and time point. A sample is eligible with at least ten cells in each compartment; a patient is paired with an eligible baseline sample and an eligible later sample (the first one). Cohorts with fewer than five paired patients were not used. Program scores are means of log_2_(CPM + 1) over program genes.

### Patient-level tests

D1: baseline-to-treatment changes were standardized within cohort; combined Spear-man correlation; two-sided P from 10,000 permutations of patients within cohort; specificity against 1,000 pairs of random gene sets, each gene replaced by one from the same expression bin of that cohort and compartment. D2: ridge regression of the on-treatment score on the baseline score of the same program (own-compartment model) or on that plus the two baseline scores of the other compartment (twin model), predictors and targets standardized within cohort on the training patients, leave-one-patient-out and leave-one-cohort-out; difference in squared error per patient tested by sign-flip permutation (10,000) with bootstrap confidence intervals; Holm correction over the two targets. D3: L2-regularized logistic regression of response on baseline scores, pooled leave-one-patient-out. Exploratory specificity controls for the remaining program pairs and for cross-sectional correlations were added after D1–D3 with a separate random-number stream, so that the pre-specified results are unchanged.

### Replication

The hypothesis (change in T-cell exhaustion markers against change in macrophage M1), the test and the criterion were written before any replication cohort was opened. The 30 series returned by the recorded searches and not examined for the discovery analysis were screened on GEO series and sample records and, where deposited, on per-cell annotation tables, without reading any expression value of a program gene. Included: GSE246613 [19], GSE286827 [20], GSE232240 [21], GSE307398 [22], GSE189926, GSE205506 and GSE264586; the reasons for excluding the other 23 are listed in the plan. For the first three, compartments follow the authors’ annotation. The last four are deposited without annotation and were typed by Leiden clustering and marker sets as above, with all marker classes (immune, epithelial, stromal and neutrophil) competing. In GSE246613 the on-treatment sample is the biopsy taken after pembrolizumab alone; in GSE264586 the cycle-2 biopsy or, when absent, the end-of-treatment biopsy; in GSE205506 untreated and anti-PD-1-treated tumour samples of the same subject. Scores, pairing, standardization, permutation and specificity control are those of D1, and the test was run once on the complete cohort set. The comparison of cohorts with and without authors’ annotation was added afterwards and is post hoc.

### Deviations from the plan

(i) The moderated *t* test, the pseudo-target calibration and the operational definition of the baselines were fixed in an addendum written after inspecting file structure but before computing effects. (ii) A first macrophage run used lanes as replication unit, contrary to the plan, and removed most guides; it was discarded and the analysis re-specified before any valid effect was computed, after seeing only that the invalid run returned no hit. (iii) No knockdown filter in macrophages (see above). (iv) The guide-assignment rule and the LPS-response program were introduced after the first valid macrophage results; the former was chosen on knockdown alone, the latter is post hoc. (v) The marker-based typing rule was corrected twice, each time on seeing a cluster-by-marker table and before any program score. (vi) The cohort inclusion rule was applied to nine series before the patient-level tests; series returned by the search but not examined at that point form the replication set. (vii) The analyses of information beyond perturbation strength (C1, C2) were specified after the marginal AUROCs of the primary test were known, and before any adjusted or matched analysis was run. (viii) For the four replication cohorts without annotation, the typing rule was amended once after reading the cluster-by-marker tables and before any program score: the cohort labelled CD45-sorted contained epithelial, stromal and neutrophil clusters that an immune-only rule assigned to macrophages or plasma cells, so all classes were made to compete and a neutrophil class was added in all four cohorts (cluster-by-marker tables of both runs in S6 Table).

## Data and code availability

All data are public: T-cell Perturb-seq [5]; T-cell screens and CRISPRa Perturb-seq, GSE174255, GSE174284, GSE190604 [6]; macrophage Perturb-seq and screens, GSE327644, GSE327124 [7]; patient cohorts under the GEO accessions listed in Methods. The ImmunoTwin code, the pre-specified analysis plan with its addenda, and all result tables are available at <u>https://w</u>ww<u>.bioinfotiget.it/gitlab/custom/immunotwin.</u>

## Author contributions

F.C. and I.M. conceived the study, performed the analyses and wrote the manuscript.

## Competing interests

The authors declare no competing interests.

## Supporting information

**S1 Text. Pre-specified analysis plan**. The plan as written before the analyses, with dated addenda A–F; each addendum states what had already been seen when it was written. Reproduced verbatim.

**S1 Table. Pre-specified criteria and outcomes**. Every criterion with the observed values and the outcome.

**S2 Table. Data sets and patient cohorts**. Perturbation data sets, functional screens, and the discovery and replication cohorts with regimen, annotation source and number of paired patients.

**S3 Table. Regulators of the T-cell effector and exhaustion-marker programs**. Donor-level effects, moderated statistics and specificity control for all targets at a false-discovery rate of 5%.

**S4 Table. Regulators of the macrophage programs**. Guide-level effects under LPS and in untreated cells for the pre-specified M1 and M2 sets and the LPS-response program.

**S5 Table. Patient-level results**. Correlations of changes for all program pairs in the discovery and replication patients at both cell-number thresholds, per cohort and combined, with permutation and specificity P values; forecast results.

**S6 Table. Cell typing of the replication cohorts deposited without annotation**. Cluster-by-marker tables of the first and of the amended run.

## Notes

### Competing Interest Statement

The authors have declared no competing interest.

