## Supplementary_Text_1 for "Causal regulators of human T-cell and macrophage activation from perturbation screens, and a replicated coupling of T-cell exhaustion markers with macrophage M1 activation in patients under checkpoint blockade"

### S1 Text. Pre-specified analysis plan

*The plan is reproduced verbatim, with its dated addenda. Its first heading follows.*

---

#### ImmunoTwin — pre-specified analysis plan

**Written 13 Nov 2026, before any result from the perturbation datasets was computed.** Timestamped by the artifact version. Every analysis in the new manuscript either follows this document or is labelled *post hoc* in the text.

Motivation: in the previous version the effector–M1 association passed its specificity control only after I switched the scoring method, having first seen the original method fail. That is a post hoc choice and it cost the claim its strength. This document fixes the choices in advance so the new results do not depend on them.

---

#### 1. Data

| Role | Dataset | Unit of replication |
|---|---|---|
| Primary perturbation map (T) | Genome-scale Perturb-seq (CRISPRi) in primary human CD4+ T cells, 4 donors × 3 conditions (Rest, Stim 8 h, Stim 48 h); donor-level DE statistics | donor (n = 4) |
| Perturbation map (macrophage) | Perturb-seq (CRISPR KO) in primary human CD14+ monocyte-derived macrophages, untreated and LPS-stimulated (GSE327644) | guide; lane as batch (single donor — see §6) |
| Functional ground truth (T) | Genome-wide CRISPRa and CRISPRi FACS screens for IL-2 and IFN-γ production in primary human T cells (Schmidt et al.; GSE174255 sgRNA counts, GSE174284 bulk RNA-seq) | sgRNA / gene |
| Orthogonal direction check | CRISPRa Perturb-seq, primary human T cells (GSE190604) | guide |
| Bulk KO re-analysis (already done) | GSE318203, GSE331345, GSE311128 | donor |
| Tumour context | ImmunoTwin atlas, 302,594 immune cells, 278 patients | patient |

#### 2. Gene programs (fixed, unchanged from the current Methods)

- **Effector**: GZMB, PRF1, IFNG, GNLY, NKG7, GZMA, FASLG
- **Exhaustion**: PDCD1, HAVCR2, LAG3, TOX, ENTPD1, LAYN, TIGIT, CTLA4
- **M1**: TNF, IL1B, IL12A, IL12B, NOS2, CXCL9, CXCL10, CD80
- **M2**: CD163, MRC1, TGFB1, IL10, VEGFA, SPP1, MSR1, MAF

Genes absent from a dataset are dropped for that dataset and the number used is reported. No gene is added, removed or reweighted after seeing a result. A program score is the mean of the per-gene z-scores across the genes present, computed within dataset and condition.

#### 3. Scoring and statistics

- Pseudobulk per replication unit (counts summed, log2 CPM+1). This is the primary scoring for every analysis, decided now, before results.
- Tests are two-sided on the unit of replication in the table above. Never on cells.
- Multiplicity: Benjamini–Hochberg within each analysis family; $q < 0.05$ is significant.
- Effect sizes with 95% CI (t-based for donor-level means, bootstrap over perturbations for ranking metrics).

- Specificity control on every program-level claim: 1,000 random gene sets matched for expression (20 quantile bins, same cells, same condition). A claim counts as specific only if the observed value exceeds the random-set null at $P < 0.05$.
- Knockdown control: perturbations whose on-target knockdown is not detectable in the data are excluded before analysis, using the efficiency table supplied with the dataset.

#### 4. The three pre-specified questions and their success criteria

**Q1 — Which genes causally move the effector and exhaustion programs?** For each perturbed gene and condition, the donor-level change in each program score versus non-targeting controls. Reported irrespective of outcome. *Success:* at least 20 genes with $q < 0.05$ on the effector program in at least one condition, and leave-one-donor-out effect correlation $r > 0.5$. If fewer, the map is reported as underpowered and the paper does not build on it.

**Q2 — Do the transcriptional effects predict independent functional outcomes?** This is the primary claim of the paper. Using the effector-program effect per gene as the only predictor, recover the hits of the independent IFN-γ and IL-2 FACS screens. *Success:* AUROC ≥ 0.65 with a bootstrap 95 % CI excluding 0.5, and above three baselines computed on the same gene set (mean expression of the gene, gene essentiality, random). Direction concordance of CRISPRa versus CRISPRi effects tested by sign test. *Failure:* reported as a negative result in the same place in the manuscript; the paper then rests on Q1 and Q3.

**Q3 — Does the perturbation map transfer to the tumour setting?** Genes whose knockdown raises the effector program are tested for enrichment among the regulators that separate effector-high from exhausted T cells in the atlas, and among the hits of published in vivo CRISPR screens on T-cell anti-tumour function. *Success:* enrichment $q < 0.05$ against the expression-matched null.

#### 5. Model benchmark (reported either way)

The T module is retrained on real perturbations and evaluated on three pre-specified splits: held-out perturbations, held-out donor, Rest ⟶ Stim transfer. Metrics: Pearson r on the perturbation deltas and direction accuracy, each against three baselines (mean perturbation effect, ridge on gene embeddings, co-expression). The outcome is reported as measured. Beating the baselines is not required for the paper; the claim stands or falls on Q1–Q3.

#### 6. Declared limitations, written before analysis

- The genome-scale T-cell data are **CD4+ cells in vitro**, whereas tumour exhaustion is predominantly a CD8+ phenotype. Any transfer to the tumour setting (Q3) is therefore an extrapolation and will be labelled as such.
- The exhaustion gene set measures surface and transcriptional markers that also rise with acute activation. Where effector and exhaustion scores move together, it will be stated that the data cannot distinguish exhaustion from activation.
- The macrophage Perturb-seq is **a single donor**, so macrophage results are descriptive: no donor-level inference, no claim of generalization across people.
- CRISPRi knockdown is not gene knockout, and CRISPRa over-expression is not physiological.
- Everything remains *in silico* with respect to tumour control: no experiment here measures killing or tumour growth.

#### 7. Stopping rule

If Q2 fails and Q1 is underpowered, the paper is not rewritten around a positive message. In that case the result is reported as a negative benchmark and the grant takes priority. No scoring method, gene set, replication unit or threshold in this document is changed in response to a result; if a deviation becomes necessary it is added to a “deviations from the pre-specified plan” section in the manuscript with its reason.

---

#### Addendum A — operational definitions fixed after inspecting file *structure* only (13 Nov 2026, before any effect was computed)

What had been seen when this was written: file sizes, column names, sample labels, counts of rows and flags. No program score, effect size, hit list or correlation had been computed.

**A1. Source of donor-level data.** The donor-stratified DE file supplied with the dataset contains estimates for *pairs* of donors (six pairs), not single donors. Donor-level effects are therefore computed here from the guide × donor × condition pseudobulk counts, as §3 already prescribes. Guides of the same target are summed within donor and condition. Targeting pseudobulks are used when flagged `keep_for_DE` by the data providers; non-targeting pseudobulks when they pass the providers' minimum-cell and total-count flags. The reference for each donor and condition is the sum of all non-targeting guides.

**A2. Test statistic for Q1.** With four donors a plain t-test has 3 degrees of freedom. The primary test is a one-sample *moderated* t-test on the four donor-level differences (empirical-Bayes shrinkage of the per-target variance towards the prior fitted across targets of the same condition and program; limma's method). The plain t-test is reported next to it. Targets must have all four donors. Hit = BH $q < 0.05$ (moderated), on-target knockdown significant in the providers' table, and target not itself a member of the program.

**A3. Calibration control for Q1.** Non-targeting guides are paired at random into pseudo-targets and passed through the identical pipeline; the fraction called as hits is reported as the empirical false-positive rate.

**A4. Leave-one-donor-out agreement (success criterion of Q1).** Primary reading, the strict one: Pearson r between one donor's effect and the mean of the other three, across *all* tested targets, averaged over the four donors; the criterion $r > 0.5$ applies to this number in the best condition for the effector program. Secondary reading, reported but not used for the criterion: the same correlation restricted to the 5 % of targets with the largest absolute mean effect in the three training donors.

**A5. Q2 ground truth and predictor.** Screen: sgRNA counts of the genome-wide CRISPRi (Dolcetto) and CRISPRa (Calabrese) FACS screens, two donors, IFN-γ and IL-2, low and high sorting bins. Per guide and donor: log2 of CPM-normalized (high + 1)/(low + 1); mean over donors; gene score = mean over guides. Gene-level P from an empirical null of 10,000 random guide sets of the same size; BH $q < 0.05$ defines hits; sign defines positive (knockdown lowers the cytokine) or negative regulators. *Primary test:* among genes tested in both experiments, recover CRISPRi IFN-γ **positive regulators** using −Δeffector at Stim 8 h. *Secondary:* negative regulators with +Δeffector; IL-2 screen; Rest and Stim 48 h; CRISPRa screen with the sign reversed; Spearman correlation between screen score and Δeffector among hits. *Baselines on the same genes:* baseline expression of the target; number of differentially expressed genes after knockdown (a generic "strong perturbation" score); essentiality proxy (depletion in the unsorted sample relative to plasmid in the same screen); random. *Robustness:* the effector score recomputed without IFNG, because the screen read-out is IFN-γ protein. *Disclosure:* the Perturb-seq and the FACS screens come from the same laboratory; they are independent experiments (different donors, libraries and read-out) but not independent groups.

---

#### Addendum B — macrophage arm: implementation corrected to match §1 (13 Nov 2026, before any valid macrophage effect was computed)

**What happened.** The first macrophage script used the *lane* as replication unit and dropped every guide with fewer than 20 cells in any single lane. With about 13 assigned cells per guide per lane this removed ~95 % of the guides: 83 (untreated) and 120 (LPS) of 584 target genes were left, with 8–17 control guides. That run returned no hit. It is not a test of anything — it deviated from §1, which states "unit: guide; lane as batch" — and no conclusion is drawn from it. What had been seen when this addendum was written: the cell and guide counts above and the line "0 hits". No effect size, gene name or direction had been looked at.

**B1. Guide assignment.** A cell is assigned to a guide when it has ≥ 3 guide UMIs and one guide accounts for ≥ 80 % of them (unchanged from the first run). The fraction of cells assigned is reported.

**B2. Unit and scoring.** Counts are summed per guide across the four lanes within condition (pseudobulk per guide, as §3 prescribes). A guide is kept with ≥ 20 cells in total. log2(CPM + 1); each gene z-scored across guides within condition; program score = mean z over the program genes present. Guide effect = score minus the mean score of the non-targeting guides. Gene effect = mean over its guides; genes need ≥ 2 guides.

**B3. Test.** Empirical null from non-targeting guides: 10,000 random sets of $n$ control guides, $n$ = number of guides of the gene (the test already used for the FACS screens, A5). Two-sided; BH within condition and program; $q < 0.05$. Genes that belong to the program are excluded from the hit list.

**B4. Controls.** (i) Calibration: non-targeting guides grouped at random into pseudo-genes of four and passed through the same test. (ii) Specificity against 1,000 expression-matched random gene sets (§3). (iii) Lane as batch: the whole analysis repeated on lanes L01+L02 and on L03+L04 separately; the correlation of gene effects between the two halves is reported as *technical* reproducibility. (iv) Stimulation check: the M1 genes must be higher in LPS than in untreated control cells.

**B5. Knockdown filter not applicable.** §3 excludes perturbations without detectable on-target knockdown "using the efficiency table supplied with the dataset". No such table exists for this dataset, and the GEO record describes the screen as CRISPR knockout in the sample titles and as CRISPR interference in the series design; a nuclease knockout does not reliably lower the target transcript. The on-target change in target-gene expression is therefore reported as a diagnostic and **not** used as a filter. This is a deviation from §3 and will be listed as such.

**B6. Criteria.** No success threshold was pre-specified for the macrophage arm, and none is added now. Whatever the number of hits, the result is reported as descriptive (single donor, §6).

---

#### Addendum C — does the effector-program effect carry information beyond perturbation strength? (14 Nov 2026, written before the analysis)

**Why.** Q2 showed that the number of differentially expressed genes after knockdown ("perturbation strength", a generic quantity) recovers the IFN-γ screen hits far better than the effector-program effect does (AUROC 0.96 vs 0.73 at Stim 8 h). The open question is whether the effector effect adds anything once strength is accounted for. The answer decides the message of the paper; it is fixed here before being computed.

**What had been seen when this was written.** The marginal AUROCs of the effector effect and of each baseline in all three conditions and both cytokines (including that at Rest the effector effect, 0.76, is above strength, 0.69); the Spearman correlations among screen hits; the CRISPRa direction check. No conditional, adjusted or matched analysis had been run.

**Data and definitions (unchanged from A5).** Genes tested in both experiments; CRISPRi screen; positive regulator = screen $q < 0.05$ with guides depleted from the high bin; non-hit = screen $q \geq 0.05$. Predictors, all standardized: $E = -\Delta\text{effector}$ (donor mean); $S = \log10(1 + \text{number of DE genes})$; $X = \log10(1 + \text{target baseline expression})$, missing set to 0; K = essentiality proxy.

**C1 — primary test.** IFN-γ screen, Stim 8 h (the condition already designated primary in A5, and the one least favourable to the effector effect). Logistic regression of positive-regulator status on S + X + K, with and without E. Likelihood-ratio test for E. *Criterion:* coefficient of E positive and LRT $P < 0.05$. Reported next to it, without a criterion: AUROC of both models in repeated stratified 5-fold cross-validation (50 repeats).

**C2 — matched comparison.** For each positive regulator, the non-hit genes within 0.2 log10 units of its S (up to the 10 nearest). Hits with fewer than 3 such controls are dropped and counted. Wilcoxon signed-rank test, across hits, of Δeffector(hit) minus the mean Δeffector of its matched controls. *Criterion:* difference negative (knockdown of a positive regulator lowers the effector program more than an equally strong knockdown of a non-hit) with $P < 0.05$.

**C3 — continuous version.** Partial Spearman correlation between screen score and Δeffector, controlling for S, X and K, across all shared genes and among screen hits (either direction). Expected sign: positive. Reported, no criterion.

**C4 — comparator program.** C1 repeated with −Δexhaustion in place of E. The exhaustion markers rise with activation, so this is not a clean negative control; it is reported to show whether any activation-linked program behaves like the effector program. No criterion.

**Secondary conditions.** C1–C3 at Rest and Stim 48 h, and for the IL-2 screen. Reported in full; they do not enter the decision.

**Decision rule.** - C1 and C2 both met ⟶ the paper may state that the effector-program effect carries functional information beyond perturbation strength, in the stimulated condition tested. - C1 met, C2 not (or the reverse) ⟶ stated as partial, with both numbers. - Neither met ⟶ the paper states that, in these data, recovery of functional hits from a transcriptional program is explained by perturbation strength, and the perturbation-strength confound is the main methodological result. The regulator maps are still reported as a resource. No predictor, transformation, caliper or condition above is changed after seeing the result.

---

#### Addendum D — patient-level test of the twin (14 Nov 2026, written before any per-timepoint score was computed)

**Why.** A digital twin has to say something about an individual patient under an intervention. Nothing in §1–§5 tests that. This addendum defines the test and what the manuscript may be called depending on its outcome.

**What had been seen when this was written.** For three public cohorts with tumour samples before and during checkpoint blockade: the list of samples, patients, time points, treatment arms and tissue sites; total cells per sample; for one cohort the authors' cluster sizes by time point; for one cohort the response label of each sample. Also known from the earlier audit: the cross-sectional exhaustion–M2 correlation in GSE169246 (r ≈ 0.01), computed by an old script that pooled blood, tumour and all time points per patient. No program score per time point, no change score and no association had been computed.

**D0. Cohorts and units.** GSE169246 (triple-negative breast cancer, paclitaxel ± atezolizumab; Zhang et al., Cancer Cell 2021 — *not* Bassez et al., as the earlier manuscript versions stated), GSE123813 (basal-cell carcinoma, anti-PD-1), GSE120575 (melanoma, anti-PD-1 and/or anti-CTLA-4; Smart-seq2). Unit = patient. Tumour-tissue samples only; blood is excluded. T compartment = all T cells; macrophage compartment = monocyte/macrophage class (authors' labels where supplied; otherwise Leiden clusters assigned by marker sets, procedure fixed in script 13). A sample is eligible with ≥ 10 cells in each compartment (the rule used throughout the manuscript); sensitivity at ≥ 30. A patient is *paired* with an eligible baseline sample and an eligible on-treatment sample (the first one when several exist; "progression" samples are not used).

**D0b. Scores.** Pseudobulk per sample and compartment, log2(CPM + 1) (log2(TPM + 1) for Smart-seq2), mean over program genes — §2 and §3 unchanged: effector and exhaustion in T cells, M1 and M2 in macrophages. Change = on-treatment − baseline. All variables are standardized within cohort before cohorts are combined.

**D1. Longitudinal coupling (primary test of the coupling).** Across paired patients: Spearman correlation between the change in the T-cell effector program and the change in the macrophage M1 program. P from 10,000 permutations of patients within cohort. Specificity: the same statistic with 1,000 pairs of expression-matched random gene sets (random on both sides), because a change in sample quality between time points would move any gene set in both compartments. *Criterion:* $r > 0$, permutation $P < 0.05$ and specificity $P < 0.05$. Reported without criterion: exhaustion–M2; each cohort separately; the two arms of GSE169246 (chemotherapy alone vs with atezolizumab); the perturbation-derived signatures of D4.

**D2. Patient-level forecast (primary test of the twin).** Target: a patient's on-treatment score, (a) macrophage M1 and (b) T-cell effector. Models, all linear, ridge penalty fixed at 1 on standardized predictors, no tuning: - *cohort mean* — the mean of the training patients; - *own compartment* — the baseline score of the same program; - *twin* — own-compartment baseline plus the two baseline programs of the other compartment. Validation: leave-one-patient-out (primary) and leave-one-cohort-out. Statistic: per-patient squared error of *twin* minus *own compartment*; sign-flip permutation test (10,000) and bootstrap 95 % CI over patients. *Criterion:* the twin has lower error than the own-compartment model with Holm-corrected $P < 0.05$ for at least one of the two targets. Both targets and both baselines are reported whatever the outcome.

**D3. Clinical response (no criterion).** Where response labels are deposited with the data (GSE120575): leave-one-patient-out logistic prediction of response from baseline samples, T-cell programs alone vs T-cell plus macrophage programs; AUROC with bootstrap CI. With about twenty baseline samples this is descriptive.

**D4. Perturbation-derived signatures (definition fixed here; derived in the next step from in-vitro data only).** *T cells:* the 50 genes most consistently decreased, at Stim 8 h, by knockdown of the IFN-γ-screen positive regulators that passed the knockdown filter (ranked by mean log2 fold change across those knockdowns; decreased in ≥ 70 % of them; the regulators themselves excluded). *Macrophages:* the 50 genes most decreased under LPS by perturbation of the canonical TLR4 pathway set of E4 (same ranking rule). Stability is reported across donors (T) and across lane halves (macrophages). These signatures enter D1 as a secondary analysis; they do not replace the §2 gene sets.

**D5. What the paper may be called.** - D2 met ⟶ "digital twin" may stand in the title, qualified as a patient-level forecast of the immune compartments. - D1 met, D2 not ⟶ the title describes coupled T-cell–macrophage changes under therapy; ImmunoTwin is named as the framework being built, and the Discussion states that it does not yet forecast. - Neither met ⟶ the manuscript is the two-compartment perturbation resource with the perturbation-strength result (Addendum C); "digital twin" is not in the title. The expected power is low: at most a few tens of paired patients across three cohorts and three platforms. A null result will be reported as inconclusive, not as evidence of absence.

---

#### Addendum E — macrophage arm: functional ground truth and data-derived programs (14 Nov 2026, written before these analyses)

**What had been seen.** Everything reported under Addendum B: the 19 hits with names and directions; split-half correlations (0.08–0.23); the fraction of control pseudo-genes with P < 0.05 (11/114 for M1); that only 22–25 % of cells carry an assignable guide; the on-target decrease (92 % of expressed targets); and the stimulation check, which shows NOS2, IL12A and IL12B close to unexpressed. Of the functional screens (GSE327124): file names, column headers and row counts only.

**E1. Functional ground truth.** FACS-sorted CRISPR-knockout screens in primary human monocyte-derived macrophages after LPS: TNF production (two donors, same guide library as the Perturb-seq) and surface CD80 (four donors, transcription-factor library). Guide score = log2 ratio of normalized counts, high over low bin, averaged over donors; gene score = mean over guides; P from 10,000 sets of control guides of the same size; BH. Positive regulator: knockout lowers the read-out, q < 0.05.

**E2. Primary macrophage test.** Among genes present in both experiments: recovery of TNF-screen positive regulators from the decrease of the M1 program under LPS, **computed without TNF** (the screen read-out is TNF protein). AUROC with bootstrap 95 % CI. *Criterion (as Q2):* AUROC ≥ 0.65 with CI excluding 0.5. Reported next to it: baselines on the same genes — perturbation strength (root-mean-square log2 fold change over the 2,000 most expressed genes), cells per gene, target expression, random; the logistic likelihood-ratio test of C1 (program effect given strength, cells and target expression); Spearman correlation and sign concordance among screen hits; the same for the CD80 screen with the M1 program computed without CD80.

**E3. LPS-response program defined from control cells.** The 50 genes with the largest LPS-versus-untreated log2 fold change in cells carrying control guides (≥ 10 CPM under LPS). Effects on this program are computed exactly as in Addendum B and reported beside the §2 M1 set, which is not replaced. This program is post hoc with respect to the stimulation check and is labelled as such.

**E4. Pathway-level test.** Canonical TLR4 ⟶ NF-κB set, fixed here from textbook knowledge: TLR4, LY96, CD14, MYD88, TIRAP, TICAM1, TICAM2, IRAK4, IRAK1, TRAF6, MAP3K7, TAB1, TAB2, IKBKB, IKBKG, CHUK, RELA, NFKB1 (members present in the library are used). Disclosure: TLR4 was already seen as a hit. Statistic: mean M1 effect of the set under LPS against 10,000 random gene sets of the same size from the library, one-sided (lower). *Criterion:* P < 0.05. Feedback inhibitors (TNFAIP3, NFKBIA, TNIP1, ZC3H12A, SOCS1, SOCS3, IL10, STAT3; members present) are tested in the opposite direction and reported without criterion; TNFAIP3 was already seen as a hit.

**E5. Guide assignment.** The 80 % dominance rule of B1 leaves three quarters of the cells unused. Alternative rules (dominant guide ≥ 60 % and ≥ 50 % of guide UMIs) are evaluated **only** on on-target knockdown: the most permissive rule whose median on-target log2 fold change is at least 90 % of that of the 80 % rule is adopted for E2–E4, and the Addendum-B results under the 80 % rule stay in the supplement. No program effect is looked at when choosing the rule.

**E6. Status of the macrophage arm.** The Perturb-seq remains a single donor and therefore descriptive; the screens add independent donors for the functional read-outs only.

---

#### Addendum D-bis — cohorts for the patient-level test, and a correction to the cell-typing rule (14 Nov 2026; still before any per-timepoint score)

**What had been seen in addition to Addendum D.** Cell-type composition per sample in the three cohorts of D0, and therefore how many patients are paired under the ≥ 10-cell rule: 11 (GSE169246), 6 (GSE123813), 1 (GSE120575; too few macrophages per Smart-seq2 sample). Eighteen patients cannot support D1 or D2. Of the additional candidate cohorts listed below: file lists and the deposited per-cell metadata tables (patient, time point, tissue, treatment, response and cell-type columns with their counts). The macrophage results of Addendum E and the gene lists of the D4 signatures had also been computed. No program score in any patient sample had been computed.

**Typing rule corrected.** The first labelling rule (arg-max of z-scores over all marker classes, non-immune ones included) mislabelled monocyte clusters as stromal and T-cell clusters as melanocytic or epithelial in CD45-sorted data; this was seen in the cluster-by-marker table. It was replaced, before any score, by the rule now documented in script 13 (immune classes only; class with the highest fraction of its own maximum; T versus NK decided by CD3D/CD3G/CD5/CD6). Where authors' annotations are deposited they are used instead.

**Cohort inclusion rule (replaces the fixed list of D0).** Every public dataset found by the GEO search recorded in `geo_twin_search.txt` is included if, from its deposited metadata alone, it has: human tumour tissue profiled by single-cell RNA-seq without sorting that removes T cells or myeloid cells; tumour samples from the same patients before and during/after a regimen containing a PD-1/PD-L1 or CTLA-4 antibody (chemotherapy-only arms are kept as a comparison arm, not pooled with them); processed counts and a patient/time-point label for each cell or sample. Candidates checked against this rule: GSE169246, GSE123813, GSE120575, GSE236581 (colorectal), GSE314906 (oesophageal), GSE301741 and GSE200996 (head and neck), GSE266919 (triple-negative breast; supersedes GSE169246 for patients present in both), GSE235863 (liver). Inclusion is decided on metadata before any score is computed, and the list of included and excluded cohorts with reasons is reported. The Bassez et al. anti-PD-1 breast cohort would qualify but its count data require a registered login on the authors' site and are not included unless the user provides them.

**Compartments in the added cohorts.** Authors' major cell-type labels are used: T cells for the T compartment; within the myeloid class, monocytes and macrophages (authors' sub-labels where they exist, otherwise the script-13 marker rule applied inside the myeloid class) for the macrophage compartment.

**Consequence for D1 and D2.** Tests, baselines, criteria and the title rule are unchanged. Primary analyses use checkpoint-containing regimens only. With cohorts on different platforms, all variables remain standardized within cohort, and leave-one-cohort-out is reported next to leave-one-patient-out. Where response is deposited (pathological or radiological), D3 is extended to those cohorts, still without a criterion.

---

#### Addendum D-ter — operational definitions for D1–D3 across the enlarged cohort set (14 Nov 2026; written while the pseudobulks were being built, before any program score in a patient sample)

**What had been seen in addition to D-bis.** For the added cohorts: the authors' cell-type and sub-type labels with their sizes, the sample × time-point tables, and (for GSE266919) the tissue site, arm and response of each sample. Sixteen of the 22 patients of GSE169246 are also in GSE266919.

- **Samples.** One sample per patient and time point: libraries or tumour regions of the same patient and time point are pooled (colorectal: several tumour regions; GSE301741: several sorted fractions — sorting changes composition, not the expression profile within a compartment). Baseline = the pre-treatment sample. On-treatment = the first later tumour sample that is eligible.
- **Cohorts used.** A cohort enters D1–D3 only with ≥ 5 paired patients, because all variables are standardized within cohort. GSE169246 is used only for patients absent from GSE266919; if fewer than 5 of them are paired it is dropped.
- **Regimens.** Primary analyses: patients whose regimen contains a PD-1/PD-L1 or CTLA-4 antibody. The chemotherapy-only arms of GSE266919 are analysed separately as a comparison and are never pooled into the primary test.

- **Scores.** log2(CPM + 1) of the sample × compartment pseudobulk, mean over the program genes detected in that cohort (§2 sets; D4 signatures as secondary).
- **D1.** Changes are z-scored within cohort; combined Spearman r; two-sided P from 10,000 within-cohort permutations of patients. Specificity: 1,000 pairs of random gene sets, each gene replaced by one drawn from the same expression bin (20 bins of mean log2 CPM over that cohort's samples, per compartment); specificity P = (1 + number of random pairs with r ≥ observed) / 1,001. Also reported, without criterion: the baseline (cross-sectional) effector–M1 correlation across the same patients.
- **D2.** Predictors and targets are z-scored within cohort using the training patients of that cohort (leave-one-patient-out), so the held-out patient does not contribute to the scaling. In leave-one-cohort-out the held-out cohort is scaled by its own baseline mean and SD. Error difference twin − own-compartment per patient; two-sided sign-flip P (10,000) and bootstrap CI (2,000); the criterion additionally requires the mean difference to be negative.
- **D3.** Responders: R (GSE314906, GSE266919; patients labelled "Out" excluded), PR or mPR (GSE301741), High or Medium pathological response (GSE200996). Logistic regression (L2, C = 1) on within-cohort z-scored baseline scores, pooled leave-one-patient-out; AUROC with bootstrap CI for T-cell programs alone and for T-cell plus macrophage programs. Descriptive.
- **Sensitivity.** Everything repeated at ≥ 30 cells per compartment.

---

#### Addendum F — outcome of D1–D2, framing chosen, and a replication test on patients not yet used (14 Nov 2026; written before any replication cohort was scored)

**Outcome of Addendum D.** D1 not met (r = 0.23, permutation P = 0.034, specificity P = 0.086, n = 90); D2 not met. By D5 the words "digital twin" do not enter the title. ImmunoTwin remains the name of the framework and of the code repository (user decision, 14 Nov 2026).

**Framing of the patient-level result in the manuscript.** "Coordinated shifts of the two compartments under checkpoint blockade, not specific to the hypothesised programs, and without forecasting value." Everything below the D1 primary line is labelled exploratory in the text: the three other program pairs, the ≥ 30-cell results, and the cross-sectional correlations at baseline and on treatment.

**Exploratory additions to the discovery set (declared here, computed after this addendum).** The random-gene-set specificity control, so far run for effector–M1, exhaustion–M2 and the perturbation-derived signatures, is also run for exhaustion–M1 and effector–M2 and for the cross-sectional baseline correlations. These are descriptive; none changes a verdict.

**F1. Replication hypothesis, fixed from the discovery set.** In the 90 discovery patients the strongest pair was Δ T-cell exhaustion vs Δ macrophage M1 (r = 0.43), chosen *after* seeing all four pairs; it therefore cannot be claimed from those patients. It is tested here on patients not used in D1–D3. - *Primary (R1):* Δ exhaustion (T) vs Δ M1 (macrophage), scores, pairing, standardization and permutation exactly as D0b/D-ter, ≥ 10 cells per compartment. **Criterion:** combined Spearman r > 0 with two-sided permutation P < 0.05. If, in addition, specificity P < 0.05 against 1,000 expression-matched random pairs, the pair is called program-specific; if not, the replication supports only the coordinated shift. - *Secondary, without criterion:* the other three pairs; R1 at ≥ 30 cells; baseline cross-sectional effector–M1. - *Minimum size.* A verdict is issued only with ≥ 25 paired patients on a checkpoint-containing regimen across the replication cohorts (power ≈ 0.6 for r = 0.43 at n = 25). Below that the result is reported as "insufficient data", not as a failure or a success. - Discovery and replication patients are never pooled for the verdict; a pooled estimate may be shown as descriptive.

**F2. Replication cohorts.** Inclusion rule of D-bis, unchanged, applied to (i) the series returned by the recorded GEO search that were *not* examined for D-bis, (ii) series returned by additional GEO queries recorded in `geo_replication_search.txt`, and (iii) the Bassez et al. breast cohort if its count data are obtained from the authors' site. Cohorts used in D1–D3 are excluded, as are GSE169246 and GSE120575 (already scored). Cohorts with < 5 paired patients are not used (D-ter). *Disclosure.* D-bis stated that every series found by the search would be included if it met the rule, but only the nine series listed there were checked against the rule before D1 was run. For two further series (GSE195832, GSE207422) only the series-level design text and file list had been retrieved (`geo_twin_candidates.txt`); their sample tables were not examined. The remaining series were never opened. No score exists for any series outside the nine. They form the replication set rather than being added retroactively to the discovery set.

**F2-bis. Replication cohorts fixed from metadata (14 Nov 2026, before any cell of these cohorts was scored).** The recorded searches (`geo_twin_search.txt`, `geo_replication_search.txt`) returned 30 series that passed a title filter and had not been examined for D-bis. Their GEO series and sample records, file lists and — for seven of them — the deposited per-cell or per-sample annotation tables were read (`geo_replication_candidates.txt`, `insp2.log–insp4.log`). What was seen: patients, time points, treatment arms, tissue, response labels, authors' cell-type labels with their sizes, file formats. No expression value of any program gene.

*Included (rule of D-bis met; expected paired patients before the cell-number filter in brackets):* - GSE246613 — triple-negative breast, pembrolizumab then pembrolizumab + radiotherapy; baseline vs the biopsy after pembrolizumab alone (the first later sample); the later post-radiotherapy biopsy is used only when the first is not eligible (D-ter). Two tumours of one patient (03T1, 03T2) are pooled. T compartment = authors' T-cell class; macrophage compartment = authors' myeloid class, sub-clusters assigned by the script-16 marker rule. [≈ 33] - GSE286827 — head and neck, durvalumab ± tremelimumab; pre vs post. T = authors' CD4+ T, CD8+ T, proliferating T and "Tstr" labels; macrophage = authors' "TAMs" label. [≈ 29] - GSE232240 — head and neck, nivolumab ± ipilimumab (IMCISION), CD45+ cells; pre vs post. T = authors' CD4 and CD8 classes; macrophage = authors' "Mono-macro" group. [≤ 18] - GSE189926 — gastric, nivolumab + disulfiram, CD45+ cells; pre vs first post. [9] - GSE205506 — mismatch-repair-deficient colorectal, anti-PD-1 ± celecoxib; untreated vs treated tumour of the same subject. [≤ 8] - GSE307398 — colorectal liver metastases, pembrolizumab + capecitabine + bevacizumab; pre vs post. [6] - GSE264586 — renal, nivolumab + ipilimumab + sitravatinib; baseline vs first later biopsy. [5]

For the last four no authors' annotation is deposited on GEO: cells are typed by the procedure of script 13 (Leiden clusters assigned by the immune marker sets), unchanged.

*F2-ter — typing of the four cohorts without annotation (14 Nov 2026; written after reading their GEO sample records and before loading any count matrix).* GSE189926 is CD45-sorted and is typed exactly as in script 13 (immune classes only). GSE205506, GSE307398 and GSE264586 are unsorted tissue; applying the immune-only rule would force tumour and stromal clusters into an immune class. For these three, **all** marker classes of script 13 (immune, epithelial, stromal) compete under the same fraction-of-maximum rule; everything else (normalization, 2,000 variable genes, 30 components, Leiden resolution 1.0, T-versus-NK decision, cells with ≥ 200 detected genes) is unchanged. The cluster-by-marker tables are saved and read before any program score is computed. In GSE205506 only tumour samples are used; untreated samples of a subject are pooled as baseline and anti-PD-1-treated samples as on-treatment. In GSE264586 the on-treatment sample is the cycle-2 biopsy, or the end-of-treatment biopsy when no cycle-2 sample exists.

*Amendment to F2-ter (14 Nov 2026; after reading the four cluster-by-marker tables and cell-class totals, before any program score in these cohorts).* The gastric data, although labelled CD45+ on GEO, contain epithelial and stromal clusters, which the immune-only rule assigned to monocytes/macrophages or plasma cells; and two large clusters with low expression of every marker set (possibly neutrophils, which the BD Rhapsody platform captures) were assigned to monocytes/macrophages for lack of a competing class. The rule is therefore made the same for all four cohorts: all marker classes compete, and a neutrophil class (CSF3R, FCGR3B, CXCR2, CMTM2) is added. All four cohorts are re-typed with this rule; the tables of the first run are kept (`*_cluster_markers_run1.csv`). No other element of the rule is changed, and clusters that remain borderline under it are left as assigned.

*Excluded, with reason:* blood or sorted/TCR-selected T cells only, hence no macrophage compartment (GSE212217, GSE315928, GSE296954, GSE280232, GSE316301, GSE317605); no pre-treatment tumour sample of the same patients, or treated and untreated samples from different patients (GSE241934, GSE279781, GSE154795, GSE193745, GSE169379, GSE320212 — its first biopsy is taken on palbociclib); fewer than five patients with both time points (GSE207422: 1; GSE288533: 2; GSE235090: 2; GSE256326: ≤ 2; GSE256118: ≤ 4; GSE131933: ≤ 4); bulk or sorted-population RNA-seq (GSE197613, GSE195832, GSE295599, GSE313101); ex vivo treatment (GSE160755).

*Pooling and units as D-ter.* Cohorts with fewer than five paired patients after the ≥ 10-cell filter drop out. All seven regimens contain a PD-1/PD-L1 or CTLA-4 antibody; three combine it with another agent (radiotherapy only after the on-treatment biopsy used; disulfiram; capecitabine + bevacizumab; sitravatinib), which is reported.

**F3. What had been seen when this was written.** All discovery results of D1–D3 (file `twin_results.json`), and — computed after the first version of this addendum, as declared above — the exploratory specificity controls of the discovery set (exhaustion–M1: $r = 0.43$, specificity $P = 0.002$). For the replication candidates: series titles and file-type lists from the recorded search, and for the Bassez cohort the data-access page. No cell, sample or score from any replication cohort.
